# Local-scale virome depiction supports significant differences between *Aedes aegypti* and *Aedes albopictus*

**DOI:** 10.1101/2022.01.13.476242

**Authors:** Arley Calle-Tobón, Juliana Pérez-Pérez, Nicolás Forero-Pineda, Omar Triana Chávez, Winston Rojas-Montoya, Guillermo Rúa-Uribe, Andrés Gómez-Palacio

## Abstract

*Aedes* spp. comprise the primary group of mosquitoes that transmit arboviruses such as dengue, Zika, and chikungunya viruses to humans, and thus these insects pose a significant burden on public health worldwide. Advancements in next-generation sequencing and metagenomics have expanded our knowledge on the richness of RNA viruses harbored by arthropods such as *Ae. aegypti* and *Ae. albopictus*; increasing evidence suggests that vectorial competence can be modified by the microbiome (comprising both bacteriome and virome) of mosquitoes present in endemic zones. Using an RNA-seq-based metataxonomic approach, this study determined the virome structure of field-caught *Ae. aegypti* and *Ae. albopictus* mosquitoes in Medellín, Colombia, a municipality with a high incidence of mosquito-transmitted arboviruses. The two species are sympatric, but their core viromes differed considerably in richness, diversity, and abundance; the viromes were dominated by a few viruses. BLAST searches of assembled contigs suggested that at least 17 virus species (16 of which are insect-specific viruses [ISVs]) infect the *Ae. aegypti* population. Dengue virus 3 was detected in one sample. In *Ae. albopictus*, up to 11 ISVs and one plant virus were detected. Therefore, the virome composition was species-specific. The bacterial endosymbiont *Wolbachia* was identified in all *Ae. albopictus* samples and in some *Ae. aegypti* samples collected after 2017. The presence of *Wolbachi*a sp. in *Ae. aegypti* was not related to significant changes in the richness, diversity, or abundance of this mosquito’s virome, although it was related to an increase in the abundance of Aedes aegypti To virus 2 (unclassified). The mitochondrial diversity of these mosquitoes suggested that the *Ae. aegypti* population underwent a change that started in the second half of 2017, which coincides with the release of *Wolbachia*-infected mosquitoes in Medellín, indicating that the population of *w*Mel-infected mosquitoes has expanded. However, additional studies are required on the dispersal speed and intergenerational stability of *w*Mel in Medellín and nearby areas as well as on the introgression of genetic variants in the native mosquito population.

## Introduction

Members of the mosquito genus *Aedes* (Diptera; Culicidae) are the primary transmitters of arboviruses such as dengue virus (DENV), Zika virus (ZIKV), and chikungunya virus (CHIKV) to humans. Two species widely distributed globally are *Ae.* (*Stegomyia*) *aegypti* (Linnaeus, 1762)—the primary vector for various arboviruses that infect millions of people in tropical and subtropical countries every year [1]—and *Ae.* (*Stegomyia*) *albopictus* (Skuse, 1885)—a native to Asia but has expanded its distribution in the last 40 years, invading not only countries in the tropical and subtropical zones but also those in the temperate zones of North America and Europe, where it has been a primary vector of some arboviruses [2,3].

Arboviruses affect several countries in the Americas, for example, Colombia; it is highly affected by DENV, ZIKV, and CHIKV [4–6]. The high incidence of arboviruses in Colombia is because of the wide distribution of *Ae. aegypti* throughout the country, where environmental and social conditions promote viral transmission [7,8]. In 1998, *Ae. albopictus* was detected in the southern part of the country, and since then, this species has expanded its distribution to numerous cities such as Medellín, where both *Ae. aegypti* and *Ae. albopictus* are infected with DENV and ZIKV [9–11]. Since 2000, the number of dengue cases in Medellín has increased continuously, with periodic outbreaks, and nearly 20,000 dengue cases have been reported in each of the last major outbreaks occurring in 2010 and 2016 [12,13]. Owing to the significant impact these mosquitoes have had on the health of this city’s residents, multiple vector surveillance and control strategies have been implemented. However, the success of these strategies has been questionable as dengue continues to occur in Medellín, periodically occurring as epidemics. The situation in Colombia reflects those in many other arboviru*s*-endemic countries, which require new approaches to control these vector-borne diseases.

Currently, there is growing scientific interest regarding the mechanism through which the mosquito microbiome and insect-specific viruses (ISVs) are involved in arbovirus transmission. Among the microorganisms that can modify the vector competence of mosquitoes, the intracellular endosymbiotic bacterium *Wolbachia* has been demonstrated *in vitro* to reduce the replication of multiple arboviruses such as DENV, ZIKV, and CHIKV in *Ae. aegypti* [14–16]; moreover, it can also alter the native host microbiome in adult mosquitoes [17].

In addition to bacteria, the microbiome of mosquitoes also includes viruses. Recent analyses of virus sequences in metagenomics data have changed our understanding of viral diversity, abundance, evolution, and roles in host biology [18,19]. In mosquitoes, pathogenic viruses represent only a fraction of the total set of viruses (virome) [20–25]. Furthermore, the identities of ISVs vary between populations and species; some ISVs are acquired from the environment, whereas others may circulate through vertical transmission, forming the virome core of mosquitoes populations [22,26]. Although the effect of many ISVs on the biology of mosquitoes remains unknown, some viruses are known to influence the mosquito immune system. For example, several ISVs may either suppress or enhance the replication of medically important arboviruses such as DENV, ZIKV, CHIKV, and West Nile viruses, suggesting that they play a role in modulating vector competence [27–29]. The insect-specific flavivirus Nhumirim virus is an example of an arbovirus suppressor as it can restrict the growth of ZIKV in mosquito cells *in vitro* and *Ae. aegypti* mosquitoes, resulting in significantly lower ZIKV infection rates in both orally infected and intrathoracically inoculated mosquitoes [30]. Although ISVs do not replicate in vertebrates, some are phylogenetically related to known pathogenic arboviruses from the families *Flaviviridae*, *Bunyaviridae*, *Rhabdoviridae*, *Reoviridae*, and *Togaviridae*. This finding has led to increasing efforts to discover and characterize more ISVs and explore ISVs as models for studying virus restriction or as potential biocontrol agents [28,29]. A recent virome description of the global populations of *Ae. aegypti* and *Ae. albopictus* reveals significant differences in the composition and diversity of ISVs found in these mosquito species. Furthermore, the abundance of some ISVs such as Phasi Charoen-like virus (PCLV) and Humaita-Tubiacanga virus (HTV) may affect the arboviruses infection capacity as well as their transmission dynamics [31].

In Medellín, Colombia, the primary vector of arboviruses is *Ae. aegypti,* but *Ae. albopictus* may potentially be a secondary vector as it can be naturally infected with DENV2 [10,32] and ZIKV [11,33]. Both Medellín and its adjacent city, Bello, sustain arboviruses at consistent rates. Thus, these cities were selected for pioneering efforts to test an alternative control strategy based on the release of *Wolbachia*-transfected *Ae. aegypti* mosquitoes, which started in 2017 [34]. These releases have not yet provided any conclusive epidemiological evidence of arboviruses control in Medellín. Furthermore, other characteristics describing the native mosquitoes’ biology such as virome composition and mosquito population diversity before and after *Wolbachia*-transfected *Ae. aegypti* release remain unexplored. Therefore, this study used a metataxonomic approach employing RNA-seq to describe the virome composition, temporal stability, and wide mitochondrial diversity of wild sympatric *Ae. aegypti* and *Ae. albopictus* captured between 2015 and 2019 in Medellín, Colombia. To our knowledge, the present study is the first to provide the estimates of the diversity and abundance of viromes in wild-caught *Ae. aegypti* and *Ae. albopictus* in Colombia. It is also the first study to report on virome diversity in a city in the Americas where *Wolbachia*-transfected *Ae. aegypti* release has been explored. This study thus provides evidence showing the local differences in the richness, diversity, and abundance of ISVs between these two mosquito species and discusses the possible impacts of the *Wolbachia*-transfected mosquitoes on the core composition of the virome and diversity of the native mosquito population on a local scale.

## Materials and Methods

### Mosquito sampling

The municipality of Medellín, Colombia, is located at 75° 34’05’’ W and 6°13’55’’ N and covers an area of approximately 376.2 km^2^. Indoor resting adult mosquitoes were captured randomly from households using mouth aspirators and entomological nets between 2016 and 2019. The captured mosquitoes were transferred alive to the Medical Entomology Laboratory, where they were killed and identified using morphological keys [35]. The blood feeding status of female mosquitoes was determined through an external examination of the abdomen. Subsequently, the mosquitoes were stored at −80°C and sorted according to the sampling periods and species. In total, 430 mosquitoes were thus divided into 14 pools.

### RNA isolation and sequencing

Total RNA was extracted from the mosquitoes using the RNeasy Mini Kit (Qiagen) according to the manufacturer’s instructions. The quality of the RNA extracts was evaluated using an Agilent 2100 Bioanalyzer (Agilent Technologies) operated by Macrogen (Seoul, Korea; www.macrogen.org). After confirming the RNA integrity number (> 7), sequencing libraries were constructed using a TruSeq total RNA library preparation kit (Illumina) after first removing the host rRNA with a Ribo-Zero-Gold (Human–Mouse–Rat) kit (Illumina). Each library was sequenced as 100-bp paired-ends on the Novaseq 6000 S4 platform (Illumina).

### Bioinformatic analysis

The paired-end reads were processed using the fastp software v.0.20.0 to remove adaptor sequences, and perform quality-based filtering [36]. The resulting high-quality reads (Phred quality score > 20) were mapped to their respective reference genomes (*Ae. aegypti* assembly AaegL5.2 [37] and *Ae. albopictus* assembly AaloF1.2 [38]) using the BWA-MEM option in VectorBase (https://vectorbase.org/vectorbase/app/) [39]. The unmapped reads were subsequently analyzed as described below.

Viral sequences were evaluated by first creating a custom database of ribosomal RNA sequences using SILVA v.132 LSU, SSU, and 5S rRNA (RF00001), 5.8S rRNA (RF00002), tRNA (RF00005), and ribonuclease P (RF00010, RF00011, and RF00373). Our custom database was then used via the software SortMeRNA v.2.1 [40] to identify, using an e-value cutoff of 10^−5^, ribosomal sequences in the unmapped reads [41]. Reads with 60% of the read length identical to ribosomal RNA sequences by >60% were excluded from further analysis. The remaining sequences were assembled using SPAdes v.3.12.0 [42], and contigs > 450 nt were classified using DIAMOND v.2.0.8 [43] and the nr viral database (updated March 2021). The analysis employed the sensitive mode for taxonomic annotation with an e-value cutoff of 10^−*5*^ and a bit score cutoff of 100 [41,44]. The output file of DIAMOND was parsed using KronaTools v.2.7.1 [45], which found the least common ancestor of the best 25 DIAMOND hits. Only contigs with identity scores of >60 % were stored. Contigs identified as viral operational taxonomic units (OTUs) were confirmed by performing BLASTn and BLASTx (https://blast.ncbi.nlm.nih.gov/Blast.cgi) searches, thus eliminating possible false positives. All OTUs were assigned to viral species based on their BLASTx identity.

### Virome diversity analysis

The number of viral reads per library was calculated by mapping the libraries against the viral contigs identified, then they were normalized by simply dividing viral reads number by the mosquito mapped reads number, and then multiplying the result by the size of the smaller sample. The alpha diversity index (i.e., diversity within each sample) was evaluated based on the observed virome richness, Shannon index, Simpson index, and evenness of each library at the virus species level using the Rhea script set [46]. Finally, the indices of the host mosquito species were compared using Kruskal–Wallis rank sum and Mann–Whitney tests employing the R software [47].

Beta diversity (i.e., viral diversity between samples) was estimated by first constructing a Bray–Curtis dissimilarity matrix [48], which considers both the shared taxonomic composition and virus abundance in the viromes. The results were plotted using nonmetric multidimensional scaling (NMDS) ordination, and the significance of the resulting clusters was tested using permutational multivariate analysis of variance (PERMANOVA; Adonis tests) employing the R software.

### Wide mitochondrial diversity between *Ae. aegypti* and *Ae. albopictus*

Paired-end reads were mapped using BWA-MEM v.0.7.17 [39,49] against the mitochondrial genomes of *Ae. aegypti* and *Ae. albopictus* (NC_035159.1 and NC_006817.1, respectively). Only reads with a mapQ score of >20 were retained using SAMtools v.1.9 [50,51]. Optical polymerase chain reaction (PCR) duplicates were removed, and read grouping was performed in Picard v.2.9.0 (http://broadinstitute.github.io/picard/). Variant calls in each pool were estimated using the “--pooled-discrete” option in the haplotype-based variant detection method implemented in freebayes v.1.2.0 [52]. Multiallelic positions at total depths of >10 x were included and indels were excluded from variant calling. To analyze sequences, we retrieved a single sequence from each sample using FastaAlternateReferenceMaker in GATK v.4.1.2.0 [53]. Then, the overall nucleotide polymorphism per site (Theta-W) was estimated using DnaSP v.6 [54], and a Kimura −2p based distance tree was constructed using MEGA X [55].

### DENV PCR assay

Because we detected DENV-3 in one of the RNA-seq samples, we performed a semi-nested reverse transcriptase (RT)-PCR to confirm this finding. The Luna Universal One-Step RT-qPCR kit was used according to manufacturer’s instructions with the primers D3_Fwd1 (5’-GACCCAGAAGGCGGTTATTT-3′) and D3-Rev1 (5’-GCCTCGAACATCTTCCCAATA-3′). These primers amplify a 1,260-bp region of the envelope gene [56]. The second PCR used the Thermo Scientific Taq DNA polymerase according to manufacturer instructions and employed the primers D3-Rev1 (5’-GCCTCGAACATCTTCCCAATA-3′) and D3-MA (5’-ACAAGCCCACGTTGGATATAG-3′). These primers amplify a 1,057-bp fragment of the envelope gene. Amplicons were purified and sequenced (Macrogen). The forward and reverse sequences were joined using BioEdit v7.2 [57], generating the consensus sequences.

### Phylogenetic analysis

The obtained consensus sequences were first aligned with reference sequences before phylogenetic analysis. The alignments were performed with MAFFT v.7.475 using the most accurate algorithm [58] and 1,000 cycles of iterative refinement. The phylogenetic tree was reconstructed from 1,000 ultrafast bootstrap maximum likelihood-based tree replicates using IQ-TREE v1.6.12 [59]. The best-fitting model was selected using ModelFinder [60]. The phylogenetic tree was drawn using FigTree v.1.4.4 (http://tree.bio.ed.ac.uk/software/figtree/).

### Ethics statement

This study focused on the *Aedes* mosquitoes and no human participants or other vertebrates were examined in the present study. Therefore, no ethical clearance was needed.

## Results

In this study, a total of 10 and 4 pooled samples of *Ae. aegypti* and *Ae. albopictus*, respectively, were analyzed using mosquitoes captured between 2015 and 2019. Each pooled sample contained 15–36 mosquitoes (Table 1), and these were subjected to RNA extraction followed by RNA sequencing. After processing the raw data, a total of 1,190,309,014 (range 93,199,032–131,645,868) 100-bp paired-end reads were generated from the 10 ribosomal RNA-depleted sequence libraries from *Ae. aegypti*; a total of 493,223,542 (range 105,740,360–131,645,868) 100-bp paired-end reads were generated from the four *Ae. albopictus* libraries. Approximately 65.3% of the reads corresponded to the mosquito sequences; therefore, a total of 405,798,910 and 191,058,189 reads were used to characterize the viromes of *Ae. aegypti* and *Ae. albopictus*, respectively, collected from Medellín, Colombia.

**Table 1.** Summary of the sequencing results showing data on pooled samples and the numbers of mosquitoes, mosquito species, read sequences, and mapped mosquito reads.

| Sample name | Mosquito species | No. of mosquitoes | Sampling time | No. of clean reads | Mosquito reads mapped |
| --- | --- | --- | --- | --- | --- |
| Ae. aeg 2015A | *Ae. aegypti* | 15 | January–May, 2015 | 93,199,032 | 88,936,067 |
| Ae. aeg 2015B | *Ae. aegypti* | 25 | September–November, 2015 | 113,389,586 | 104,933,849 |
| Ae. aeg 2016A | *Ae. aegypti* | 35 | February–June, 2016 | 129,886,418 | 88,083,776 |
| Ae. aeg 2016B | *Ae. aegypti* | 33 | July–November, 2016 | 122,388,802 | 74,256,344 |
| Ae. aeg 2017A | *Ae. aegypti* | 31 | March–June, 2017 | 105,740,360 | 56,373,655 |
| Ae. aeg 2017B | *Ae. aegypti* | 34 | July–December, 2017 | 128,342,842 | 75,574,534 |
| Ae. aeg 2018A | *Ae. aegypti* | 35 | March–June, 2018 | 119,858,346 | 72,780,110 |
| Ae. aeg 2018B | *Ae. aegypti* | 33 | July–November, 2018 | 131,645,868 | 85,013,726 |
| Ae. aeg 2019A | *Ae. aegypti* | 31 | January–June, 2019 | 123,578,244 | 74,904,614 |
| Ae. aeg 2019B | *Ae. aegypti* | 28 | July–November, 2019 | 122,279,516 | 76,372,131 |
| Ae. alb 2017 | *Ae. albopictus* | 25 | August–December, 2017 | 124,650,502 | 81,419,768 |
| Ae. alb 2018 | *Ae. albopictus* | 36 | March–August, 2018 | 114,442,616 | 63,806,581 |
| Ae. alb 2019A | *Ae. albopictus* | 34 | January–June, 2019 | 130,821,410 | 71,501,186 |
| Ae. alb 2019B | *Ae. albopictus* | 35 | July–November, 2019 | 123,309,014 | 85,437,818 |

### Virome characterization

After annotating and analyzing the contigs with DIAMOND BLASTx annotation and KronaTools, respectively, the contigs were verified; 125 of them were identified as viral OTUs (86 from *Ae. aegypti* and 39 from *Ae. albopictus*) comprising 21 virus species. These virus species were related to ISVs, arboviruses, and plant viruses belonging to nine different viral families: *Flaviviridae, Totiviridae, Reoviridae, Phenuiviridae, Iflaviridae, Bromoviridae, Metaviridae, Xinmoviridae, and Orthomyxoviridae*; however, six viruses remained unclassified.

Sequences from each library were mapped to the identified viral contigs to determine the abundance of each identified virus species. The rarefaction curves shown in Fig 1A are based on the number of viral sequences; these show that the curves describing samples are approaching the saturation plateau, suggesting that most viruses were detected in this experiment. In addition, *Ae. albopictus* curves showed a tendency to plateau faster, suggesting that the virome of this species is less diverse than that of *Ae. aegypti*. Furthermore, significantly lower numbers of viral sequences were detected in *Ae. albopictus* (Fig 1B).

**Figure 1.**
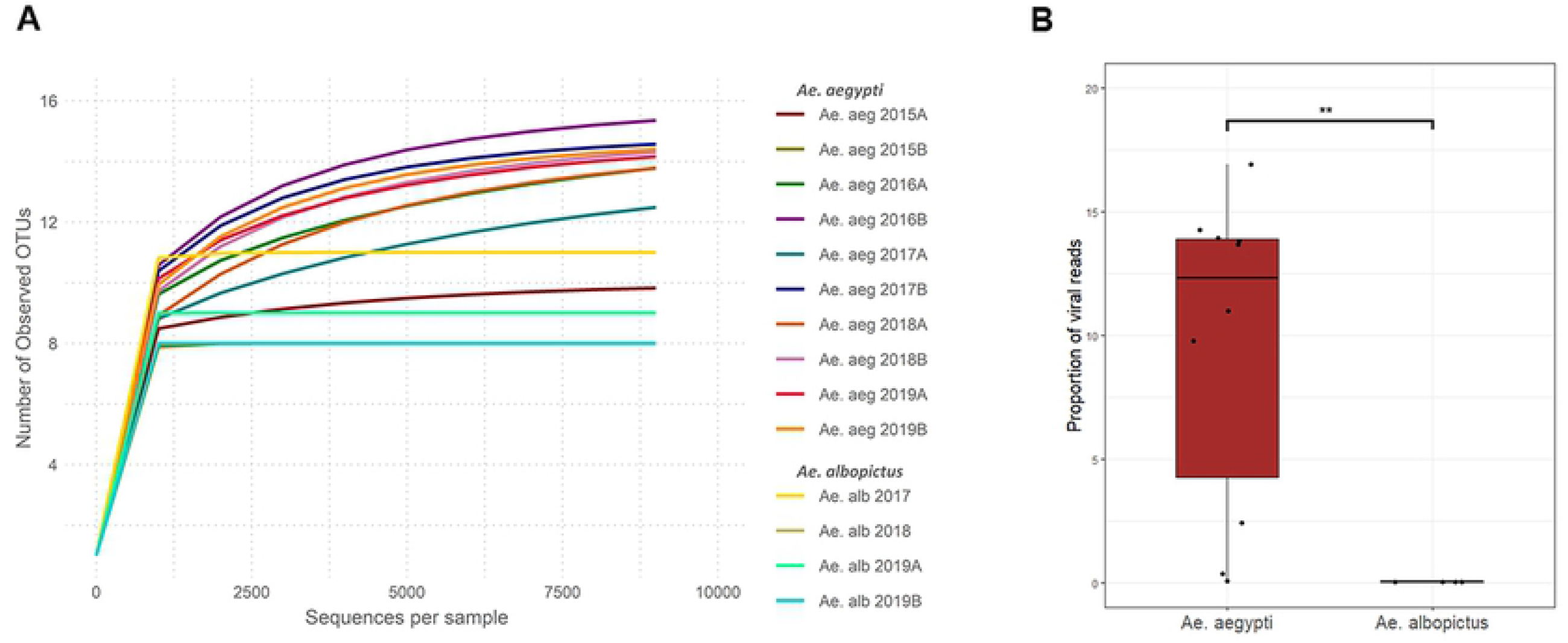
**A.** Rarefaction curves representing the viruses in *Ae. aegypti* and *Ae. albopictus*. **B.** Proportion of viral sequences relative to the number of mosquito host sequences. Mann–Whitney Test: *p* < 0.05 (*) and *p* < 0.01 (**)

A total of 75,219,419 viral sequences (6.3% of libraries) were identified in *Ae. aegypti*, which were assigned to 17 viral species. The virome of this species was dominated by four viruses, which accounted for approximately 90% of the viral sequences. Of the total viral sequences, 52% corresponded to PCLV (*Phenuiviridae*); PCLV was present in all libraries and was the most abundant virus in almost all samples, except for Ae. aeg 2015B, where Aedes anphevirus (AaAV; *Xinmoviridae*) was the most abundant virus detected (Fig 2). The second and third most abundant viruses were Kwale mosquito virus (21.5%) and Guadeloupe mosquito virus (GMV; 12.2%), respectively; both viruses are unclassified but are probably mosquito-specific viruses. Finally, cell fusing agent virus (CFAV; *Flaviviridae*) accounted for 3.7% of the total viral sequences.

**Figure 2.**
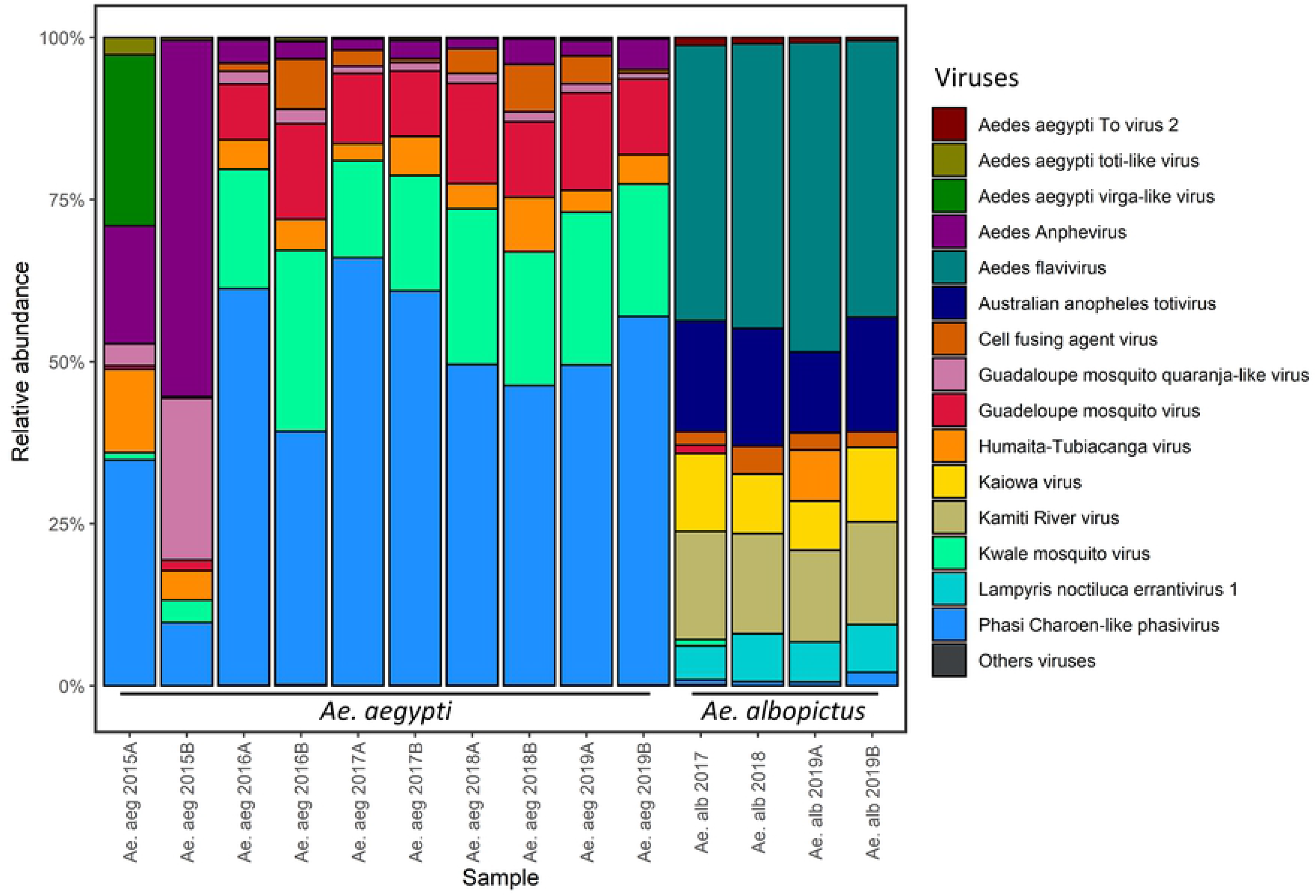
Relative abundance of virus species in all samples.

Interestingly, DENV3 (*Flaviviridae*) was detected in one of the samples from 2016. We confirmed the presence of this virus through RT-PCR targeting a fragment of the envelope gene. Subsequent phylogenetic analysis identified this sequence as DENV3 genotype III (**Fig S1**). No other arboviruses were identified in this study. The *Ae. aegypti* virome composition was fairly conserved among the samples throughout the 5 years of sampling, suggesting that the virome of *Ae. aegypti* in Medellín is stable over time.

The virome of *Ae. albopictus* had less viral sequences and species than that of *Ae. aegypti* (Fig 1). In *Ae. Albopictus,* a total of 12 viral species were identified, which were pooled into five families (84,339 reads, 0.02% of the total). Aedes flavivirus (AeFV; *Flaviviridae*) was the most abundant virus, accounting for 44.4% of the viral sequences in all samples (Fig 2). Australian Anopheles totivirus (*Totiviridae*) accounted for 16% of viral sequences, and Katimi river virus (*Flaviviridae*) accounted for 15.3%. PCLV was predominant in the virome of *Ae. aegypti*, whereas it accounted for only 1% of the total viral sequence in *Ae. albopictus*. Of all viruses detected in *Ae. albopictus*, only eight were present in multiple samples (Fig 3). Tobacco streak virus (*Bromoviridae*)—a plant virus—and Kwale mosquito virus were detected in sample Ae. alb 2017, whereas GMV (unclassified viruses) and HTV were detected in sample Ae. alb 2019A. These results suggest that *Ae. albopictus* acquired these viruses from the environment.

**Figure 3.**
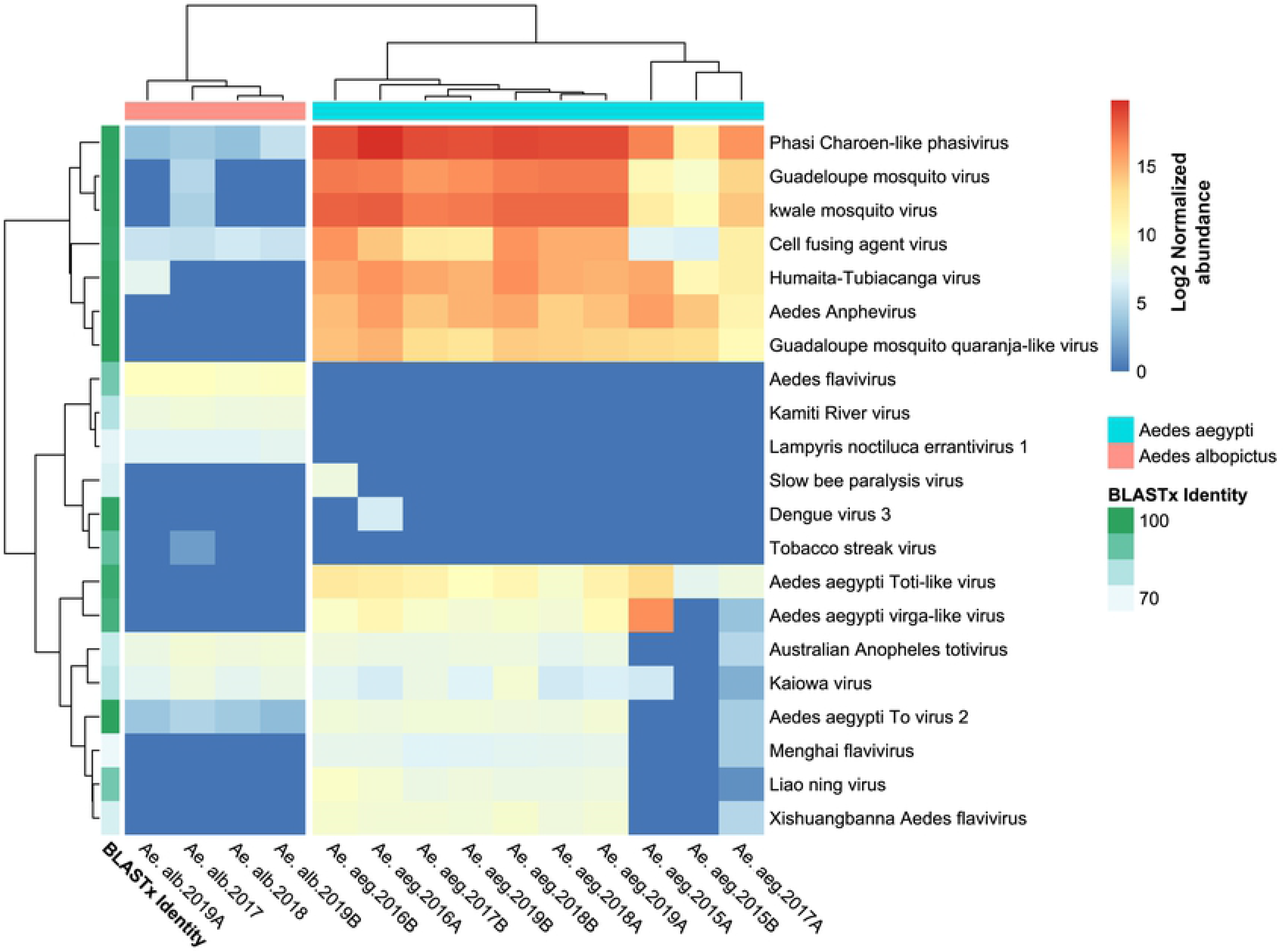
Heatmap showing the normalized abundance of reads on a log2 scale. The viral species names are from the taxonomic annotation based on DIAMOND and KronaTools. The average BLASTx identity of each virus species to a reference sequence is shown in the blue-green boxes on the left.

### Virome comparison between the *Aedes* spp

The two *Aedes* spp. differed in the composition and abundance of virus species in their respective viromes. Fig 3 clearly shows the hierarchical clustering of virus species in each mosquito species based on the Euclidean distance matrix. In total, 21 virus species were identified, of which only eight were shared between the mosquito species studied (Australian Anopheles totivirus, CFAV, PCLV, GMV, HTV, Kwale mosquito virus, Kaiowa virus, and Aedes aegypti To virus 2). Similarly, viruses in both mosquito species had significantly different levels of alpha diversity, as evident from the indexes of diversity and richness (Fig 4A). Although the *Ae. aegypti* virome is richer in virus species, the *Ae. albopictus* virome is more diverse, based on Shannon and Simpson indices. This higher diversity may be because of the higher evenness of the *Ae. albopictus* viral community.

**Figure 4.**
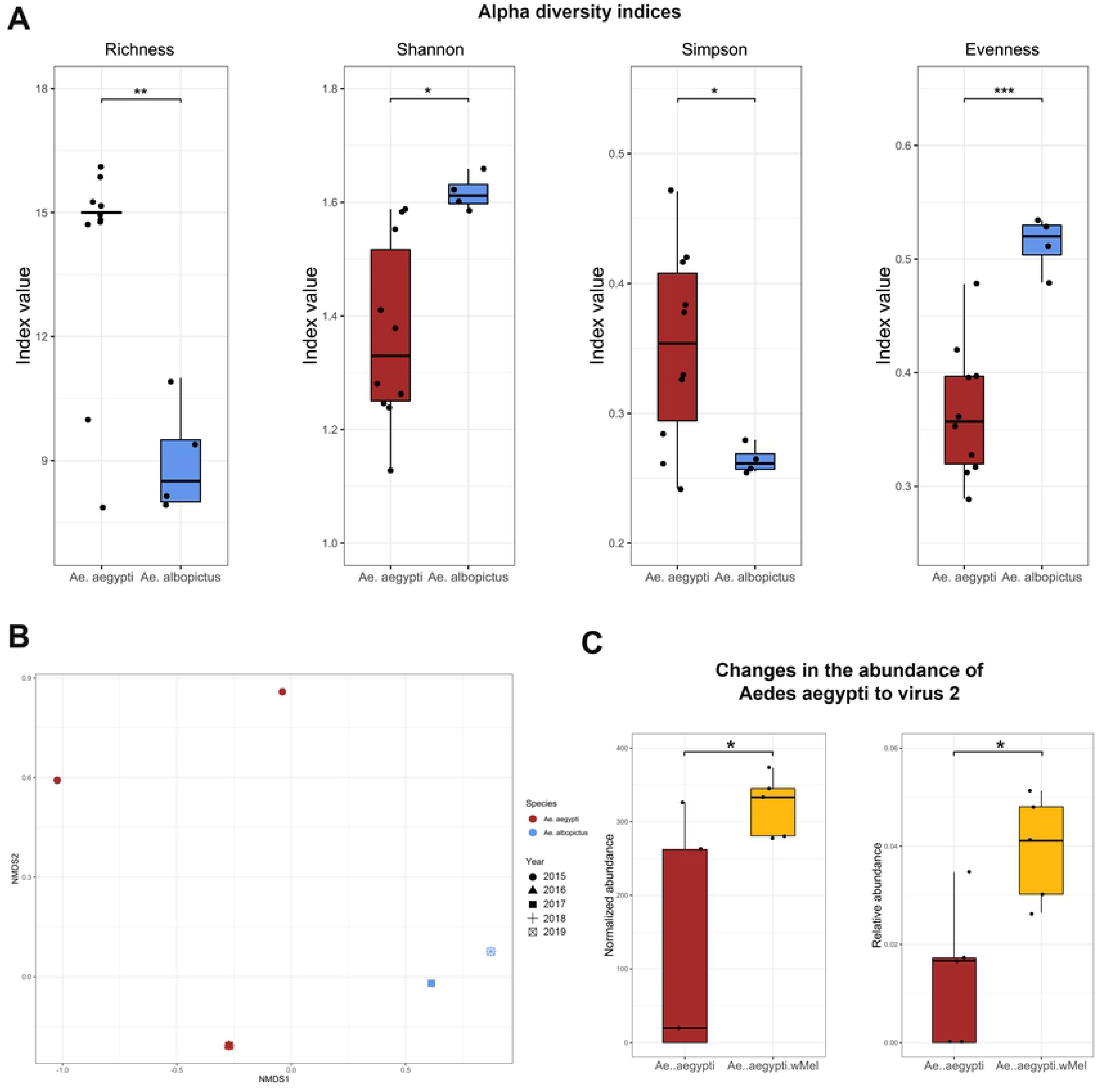
Diversity analysis of the *Ae. aegypti* and *Ae. albopictus* viromes. **A.** Alpha diversity of viruses in *Ae. aegypti* and *Ae. albopictus* at the species level. Mann–Whitney Test: *p* < 0.05 (*), *p* < 0.01 (**), and *p* < 0.001 (***). **B.** Nonmetric multidimensional scaling (NMDS) of viruses associated with the two mosquito species. STRESS = 0.023, PERMANOVA test on mosquito species: p = 0.004 and R2 = 0.59. **C.**Comparison of normalized and relative abundance levels of Aedes aegypti To virus 2 in the samples of *Ae. aegypti* with and without *Wolbachia* sp.

Beta diversity is based on Bray–Curtis dissimilarities that were calculated from the normalized abundance of viral species and then analyzed via unconstrained ordination using the PERMANOVA test on mosquito species (p = 0.004 and R2 = 0.59) and NMDS. Fig 4B shows a clear separation of viral communities according to mosquito species, suggesting that the viromes of these two mosquito species differ significantly in their compositions.

### Effects of *Wolbachia* on the *Ae. aegypti* and *Ae. albopictus* viromes

The presence of the intracellular symbiotic bacterium, *Wolbachia*, was detected in all *Ae. albopictus* samples, which might have been because this species is naturally infected by the bacterium. In contrast, *Ae. aegypti* is not a natural host for *Wolbachia;* however, *Ae. aegypti* infected with *Wolbachia* have been released in Medellín since 2017. *Wolbachia* sequences were detected in five *Ae. aegypti* samples that were collected from mid-2017 to 2019. The consensus contigs of the gene encoding the surface protein (*wsp*) were used for phylogenetic analysis (Fig 5), which shows that *Ae. albopictus* was infected by *Wolbachia* spp. from clades A and B. In *Ae. aegypti*, *Wolbachia* sp. belonging to clade A, which is related to *Drosophila melanogaster* isolates, was detected. This result was expected because the released mosquitoes were transfected with the *wMel* strain.

**Figure 5.**
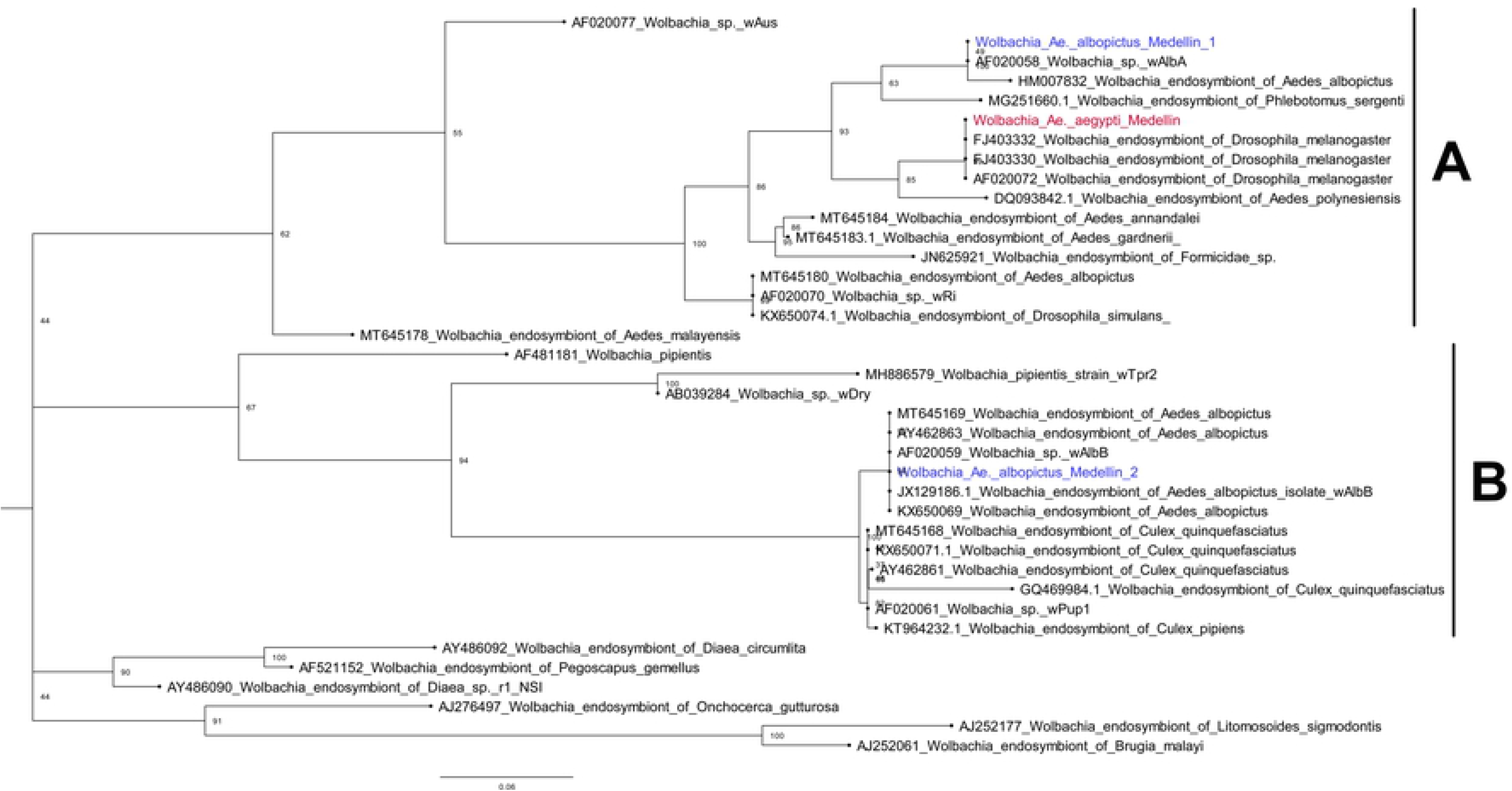
Phylogenetic reconstruction of the *Wolbachia* consensus sequences associated with *Ae. albopictus* (blue) and *Ae. aegypti* (red).

Whether the *Wolbachia* infection in *Ae. aegypti* could influence the richness or diversity of the *Ae. aegypti* virome was investigated. For this purpose, *Ae. aegypti* samples with and without *Wolbachia* sp. were compared. No significant differences in richness or diversity were found between the samples; however, Aedes aegypti To virus 2, an unclassified RNA virus, was significantly more abundant in samples with *Wolbachia* (Fig 4C).

### Mitochondrial diversity of *Aedes mosquitoes*

On average, 459,117 (± 222,695) reads were mapped in *Ae. aegypti*, whereas 443,887 (± 195,269) reads were mapped in *Ae. albopictus* (Table 2). A total of 285 and 150 single-nucleotide polymorphisms (SNPs) were detected in *Ae. aegypti* and *Ae. albopictus*, respectively (Table 2).

**Table 2.**
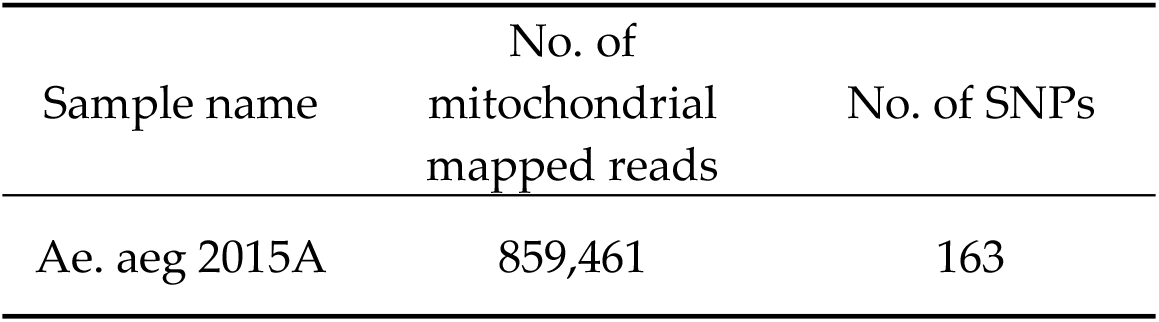
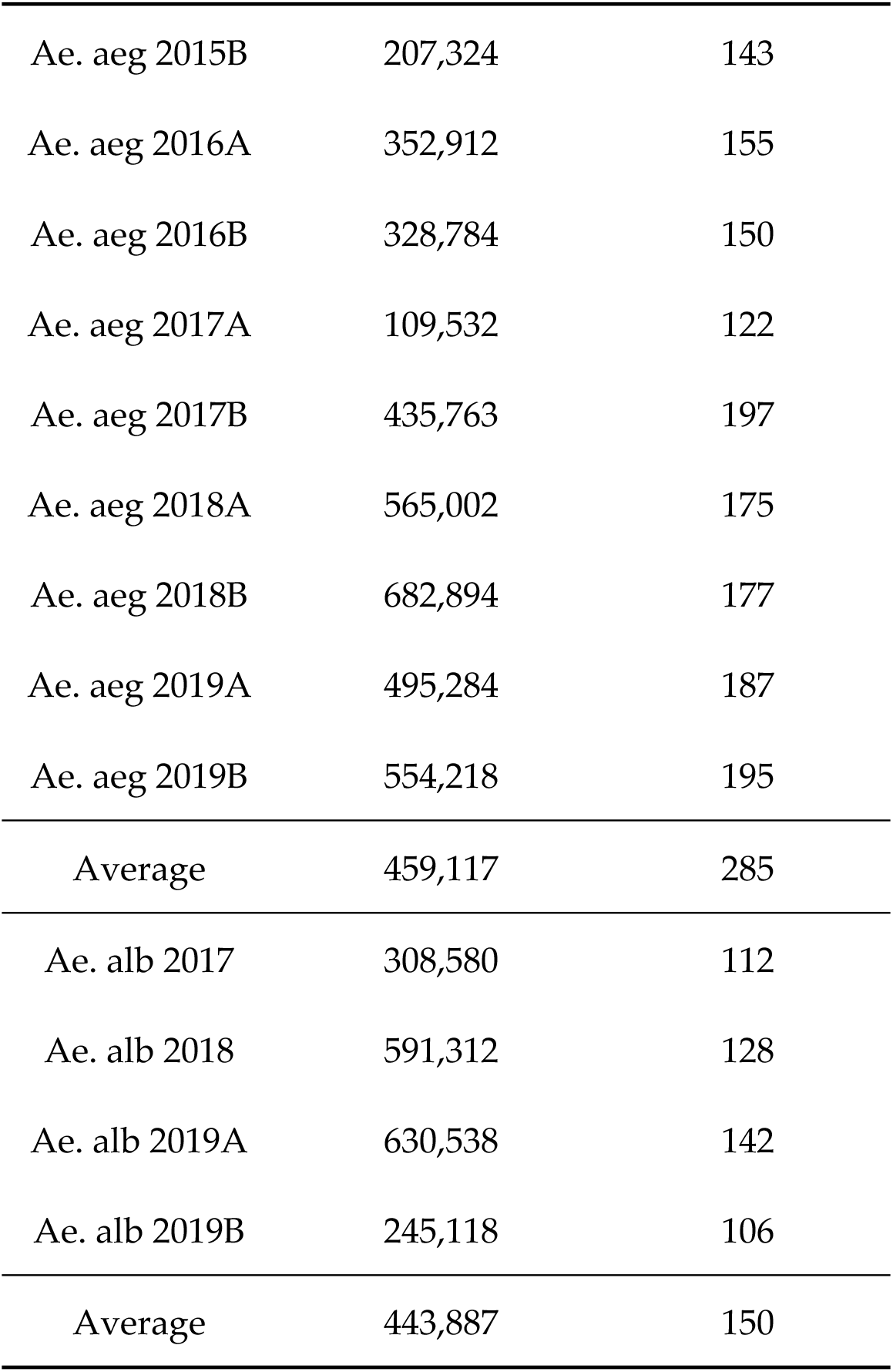
Results of mitochondrial diversity analysis showing the numbers of mitochondrial mapped reads and SNPs.

| Sample name | No. of mitochondrial mapped reads | No. of SNPs |
| --- | --- | --- |
| Ae. aeg 2015A | 859,461 | 163 |
| Ae. aeg 2015B | 207,324 | 143 |
| Ae. aeg 2016A | 352,912 | 155 |
| Ae. aeg 2016B | 328,784 | 150 |
| Ae. aeg 2017A | 109,532 | 122 |
| Ae. aeg 2017B | 435,763 | 197 |
| Ae. aeg 2018A | 565,002 | 175 |
| Ae. aeg 2018B | 682,894 | 177 |
| Ae. aeg 2019A | 495,284 | 187 |
| Ae. aeg 2019B | 554,218 | 195 |
| Average | 459,117 | 285 |
| Ae. alb 2017 | 308,580 | 112 |
| Ae. alb 2018 | 591,312 | 128 |
| Ae. alb 2019A | 630,538 | 142 |
| Ae. alb 2019B | 245,118 | 106 |
| Average | 443,887 | 150 |

In general, the number of SNPs per sample was higher in *Ae. aegypti* than in *Ae. albopictus,* except for sample Ae. aeg 2015A (Fig 6). However, variable sites were observed throughout the entire mitochondrial genome in both species. SNP density peaked at around position 7,000 of the mitochondrial genome, a region involved with coding for the ND5 gene in both species (Fig 7). In addition, a second high-density SNP region located between positions 12,400 and 14,600 of the mitochondrial genome was observed in *Ae. aegypti* samples. This location harbors coding regions of the ND1, tRNA-Leu, tRNA-Val, and 16s, and 12s rRNA genes (Fig 7).

**Figure 6.**
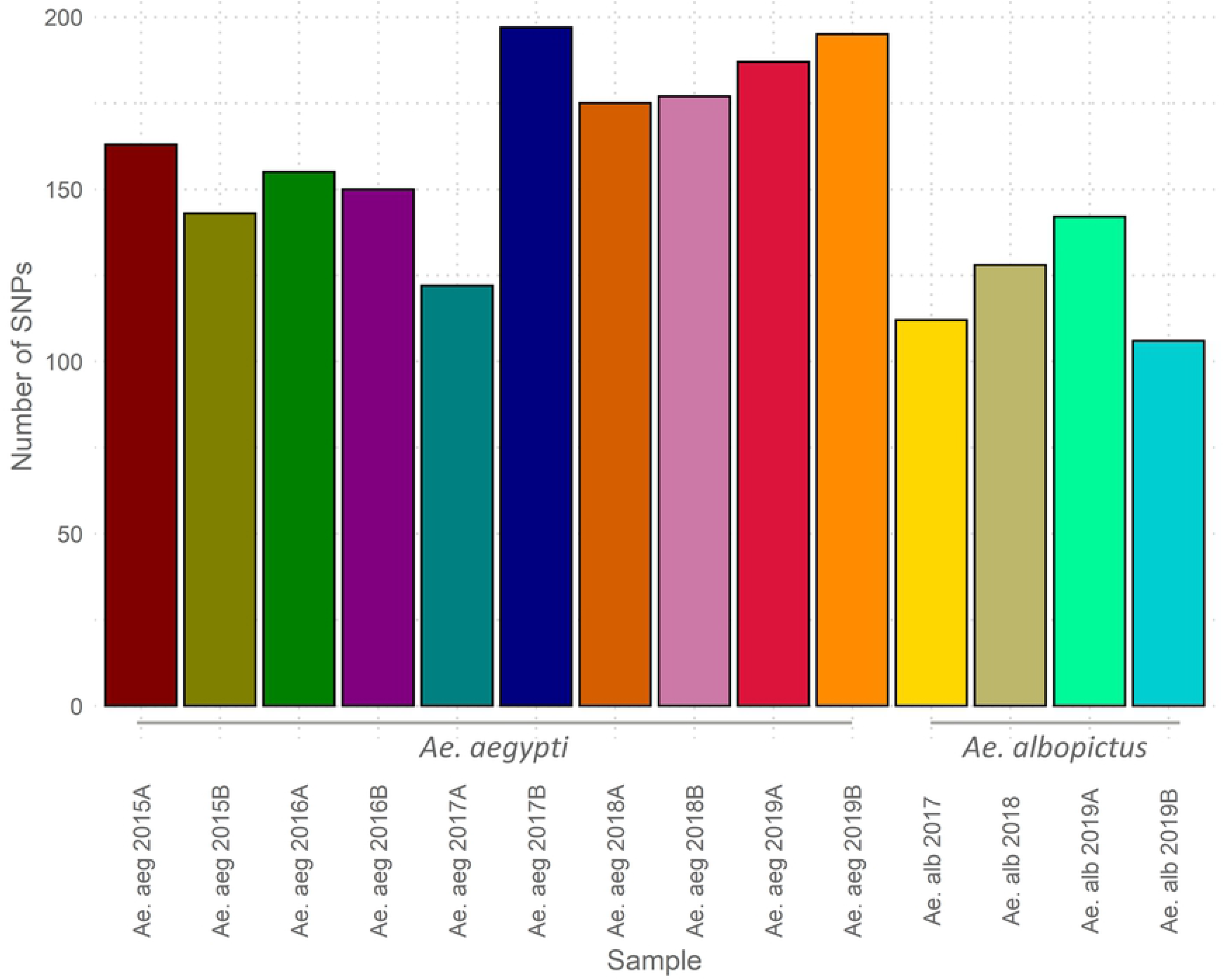
Number of mitochondrial SNPs in *Ae. aegypti* and *Ae. albopictus* samples collected between 2015 and 2019 in Medellín, Colombia.

**Figure 7.**
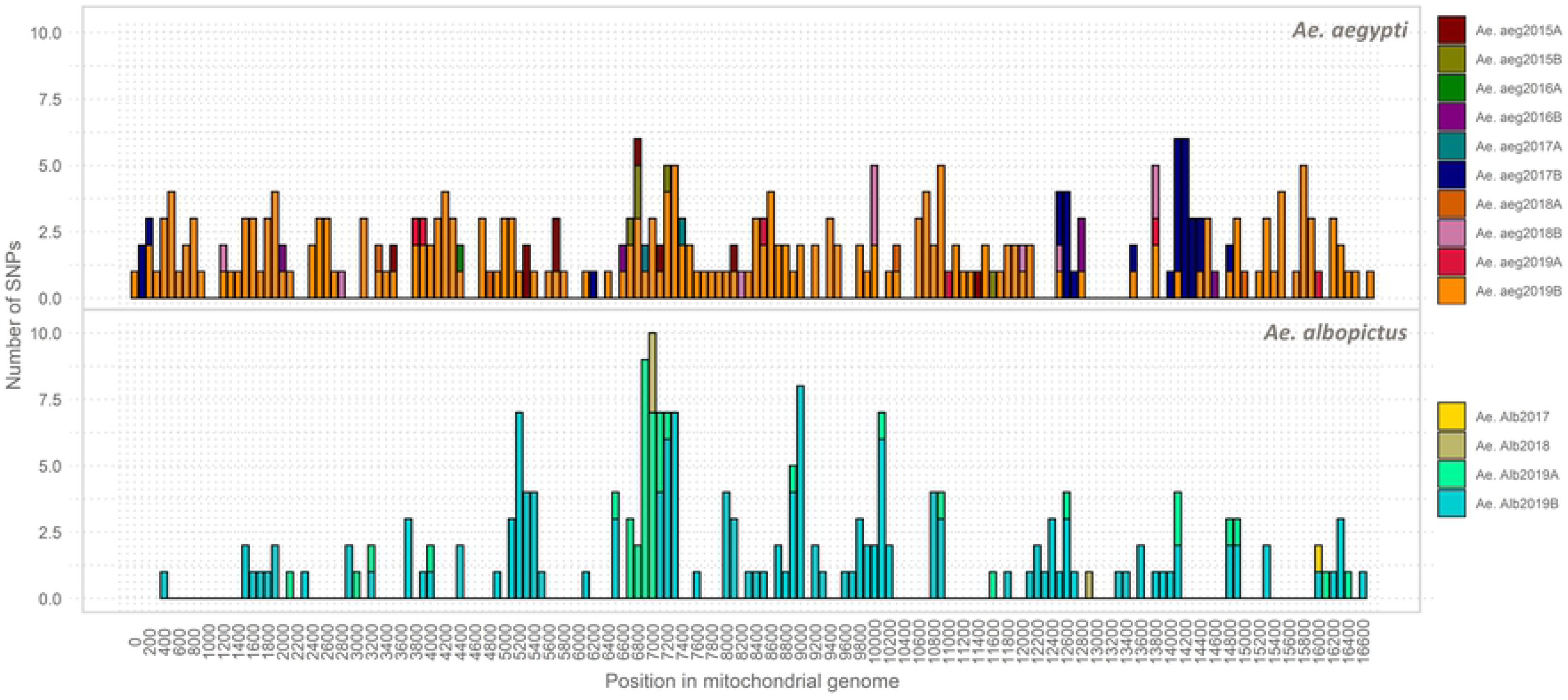
Mitochondrial SNP density per site in *Ae. aegypti* and *Ae. albopictus* samples collected between 2015 and 2019 in Medellín, Colombia.

The overall nucleotide polymorphism (Theta-W) was 0.004 for *Ae. aegypti*, whereas it was 0.002 for *Ae. albopictus*. Thus, the neighbor-joining tree showed two well-supported clades harboring both species. A secondary clade including samples collected after June 2017 (i.e., sample Ae. aeg 2017A) was also revealed in *Ae. aegypti* (Fig 8).

**Figure 8.**
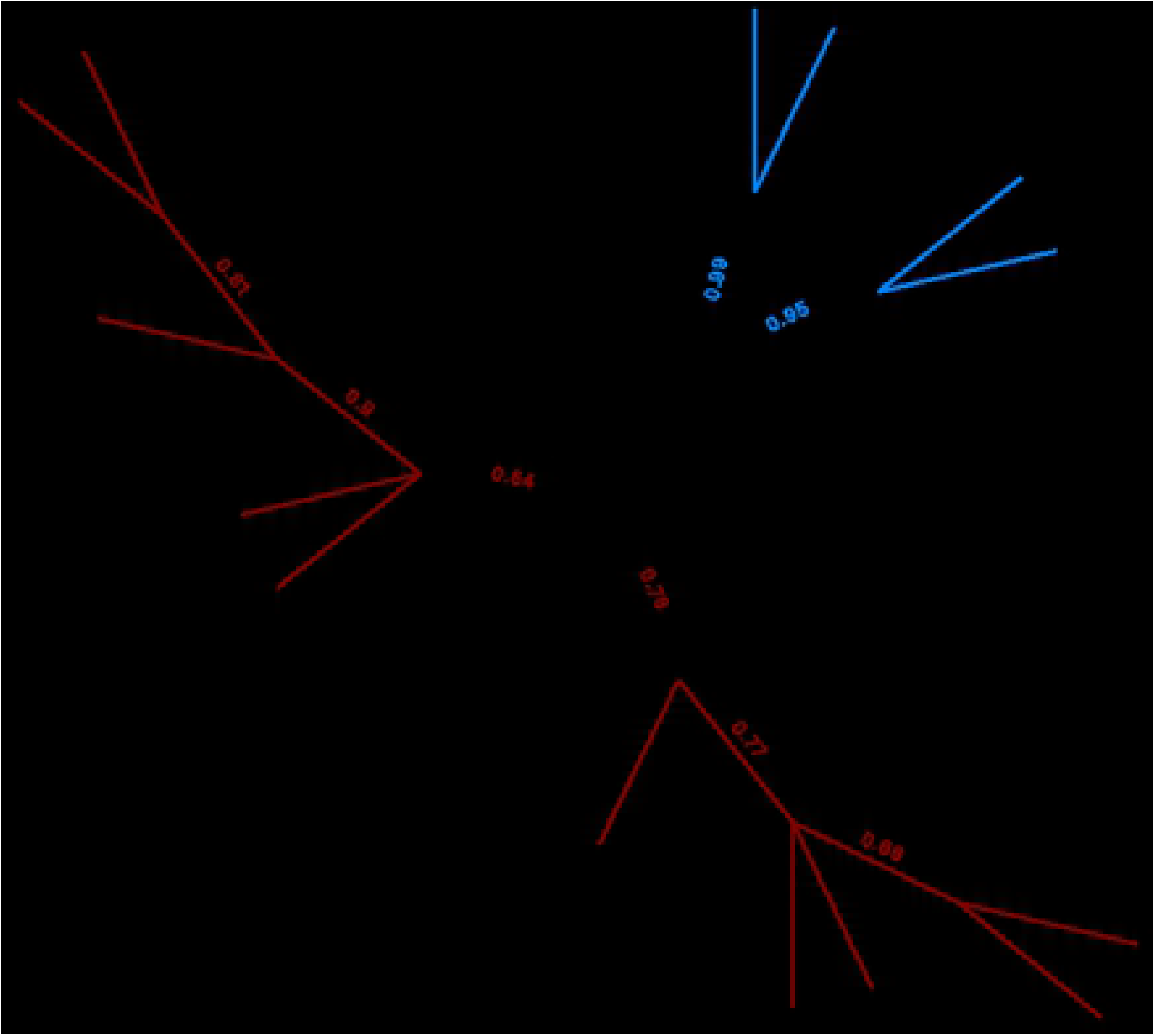
Distance-based tree based on mitochondrial sequences in *Ae. aegypti* and *Ae. albopictus* collected between 2015 and 2019 in Medellín, Colombia.

## Discussion

Recent studies on mosquito microbiomes have begun to elucidate the complex interactions between microorganisms and their hosts as well as the effects the former has on the latter. Most studies have focused on the bacteriome, thus the role and transmission of the virome, which comprises a highly diverse community of viruses, are less understood. Metagenomic studies rely on database analysis, which is a problem in virome studies because sequence information for many viral families and genera is still limited. Therefore, this study used stringent restrictions such as OTUs identity and cover greater than 60% to reduce the possibility of reporting false positives. Although false positive errors are minimized through this approach, it can also result in the underestimation of the true diversity of viruses in these mosquito populations.

This study, to the best of our knowledge, is the first to characterize the viromes of *Ae. aegypti* and *Ae. albopictus* populations in Colombia, where mosquito-borne diseases such as dengue, Zika, and chikungunya are major public health concerns [5]. The findings of the present study add significant information to the existing knowledge regarding the mosquito viromes in the Americas and the world. The study results reveal that the compositions of the viromes of sympatric, field-caught *Ae. aegypti* and *Ae. albopictus* differ significantly, suggesting that virome composition is species-specific and reflects differences in the host evolutionary history, host immunological response, virus– mosquito interactions, and perhaps vector competence. The two mosquito species also differ largely in the richness of virus species associated with them and the abundances of these viruses. This result was consistent across all comparisons (Fig 2).

A larger proportion of viral sequences were associated with *Ae. aegypti*; these sequences represented a greater richness of viruses than those associated with *Ae. albopictus*. These results support the hypothesis that *Ae. aegypti* has a higher capacity to carry viruses and, therefore, to disperse these viruses more widely. This also suggests that this mosquito species is more susceptible to infection by arboviruses such as DENV, ZIKV, and CHIKV [31]. Globally, the *Ae. aegypti* virome is highly variable in terms of the richness of viral species [22,31]. In the present study, the *Ae.* aegypti virome comprised least 17 virus species predominated by PCLV, which is consistent with the results of previous studies showing that this virus superinfects *Ae.* aegypti, for example, PCLV has been reported to be a predominant ISV in *Ae. aegypti* populations in Australia [41], Guadeloupe [61], South China [62], Thailand [63], Brazil [31], and Grenada [64]. Interestingly, PCLV is less abundant in African populations such as those in Senegal, and it is absent in Gabon, where the dominant subspecies is related to *Ae. aegypti formosus*, which is considered less susceptible to arbovirus infection [65].

GMV and Kwale mosquito virus were also highly abundant in *Ae. aegypti* samples. GMV has been identified recently in *Ae. aegypti* from Guadeloupe, where it shows high abundance in the mosquito virome [61,66]. Kwale mosquito virus was detected in a metagenomic analysis of *Ae. aegypti* mosquitoes from Kenya; it is phylogenetically closely related to Hubei mosquito virus 2 and GMV related viruses [22]. GMV and Kwale mosquito virus are currently considered unclassified viruses, and their effects on the biology of *Aedes* remain unknown. Another abundant member of the *Ae. aegypti* virome is CFAV, a flavivirus that was the first insect-specific flavivirus found in *Aedes* [67]. Since then, it has been detected in multiple regions [22,24,68–70]. Furthermore, CFAV infection can significantly enhance DENV replication in *Ae. aegypti* cells [71].

In this study, *Flaviviridae* was the most diverse family, and three ISVs as well as one arbovirus (CFAV, Menghai flavivirus, Xishuangbanna AeFV, and DENV3) were detected in most samples. Menghai flavivirus has been isolated from *Ae. albopictus* mosquitoes in China, and it is phylogenetically related to Xishuangbanna AeFV, which belongs to the *Aedes*-associated flaviviruses cluster [72]. Although ISVs are abundant and highly prevalent, arbovirus infections in wild mosquitoes are rare. Nonetheless, DENV3 was detected in sample Ae. aeg. 2016A. Results of the phylogenetic analysis revealed that this virus belongs to genotype III, the same genotype that has been circulating in Colombia and other South American countries [56]. A dengue epidemic occurred in Medellín in 2016 [13]; however, the *Ae. aegypti* virome from that year did not differ from those from other years.

ISVs from the *Totiviridae* family (Aedes aegypti toti-like virus and Australian Anopheles totivirus) and one from the *Iflaviridae* family (iflavirus) were detected in sample Ae. aeg 2016B. The latter is related the slow bee paralysis virus (72.2% BLASTx identity), although it may also be a new noncharacterized *iflavirus*. Similarly, several viruses from *Aedes* mosquitoes have been tentatively identified as Iflavirus-like viruses [66,73,74]. These results indicate that this virus family needs to be expanded and revised. Furthermore, AaAV (*Xinmoviridae*) was detected in all samples. This *Ae. aegypti-*associated virus has been detected in field-caught and colony mosquitoes as well as in cells lines. AaAV can interact with *Wolbachia* and, in the process, influence DENV replication in cell lines [75].

Virus species richness was lower in *Ae. albopictus* samples, where 12 virus species were detected. The core virome of *Ae. albopictus* mainly comprised eight viruses that were found in multiple samples. The most diverse set of viruses belonged to the *Flaviviridae* family and included three ISVs, namely, AeFV, CFAV, and Kamiti river virus. AeFV is a flavivirus associated with both *Ae. albopictus* and *Ae. flavopictus* [76]. A temporal study of AeFV in *Ae. albopictus* in China suggests that it is a seasonal virus [77]. Because Colombia is a tropical country, AeFV may be perennially present in *Ae. albopictus*.

Australian Anopheles totivirus was the second most abundant species in *Ae. albopictus,* and it was present in all samples in addition to Kaiwá virus, Aedes aegypti To virus 2, Lampyris noctiluca errantivirus 1 (*Metaviridae*), and PCLV. In contrast to the findings obtained from *Ae. aegypti*, PCLV was rare in *Ae. albopictus samples*. Notably, both PCLV and CFAV have been reported to inhibit ZIKV, DENV, and La Crosse virus (*Peribunyaviridae*) in the *Ae. albopictus* cell line Aa23 [78].

More unique viruses per sample were detected in *Ae. albopictus* than in *Ae. aegypti*; furthermore, Tobacco streak virus (*Bromoviridae*), HTV, Kwale mosquito virus, and GMV were each detected in only one sample, suggesting that these viruses have been acquired from the environment. The higher level of variability in the virome composition of *Ae. albopictus* may be because of this mosquito’s tendencies to breed in natural places and feed on different types of vertebrates [79,80], thus exposing itself to more viruses in the environment.

Mosquitoes are not known to be vectors of plant viruses; however, they can acquire these viruses from nectar during feeding; thus, some plant viruses have been detected in mosquito viromes [24,25]. For example, in China, a plant-specific Tymoviridae-like virus that was isolated from *Culex* spp. was able to replicate in mosquito cells, suggesting that it had the potential to be transmitted by mosquitoes [81].

Despite the lower level of virus richness in *Ae. albopictus* than that in *Ae. aegypti*, the Alpha diversity of the *Ae. albopictus* virome was higher. This is because the abundance levels of viruses in *Ae. albopictus* were more evenly distributed. Multiple virus sequences identified in this study are consistent with other recently described sequences that currently lack a formal taxonomic classification. Therefore, the mosquito virome needs to be better characterized and considered in relationship with vector competence. Furthermore, the mosquito virome diversity can be easily influenced by leading mosquitoes to establish colonies [22].

An important difference between *Ae. albopictus* and *Ae. aegypti* is the naturally high prevalence of the intracellular bacterium *Wolbachia* in the former [82,83] and its absence in the latter [84]. *Wolbachia* colonization in *Cx. quinquefasciatus* reduces the abundance and richness of its associated viruses, which may also explain the differences between the *Ae. aegypti* and *Cx. quinquefasciatus* viromes [61]. The relative abundance levels of the resident bacteria in adult *Ae. aegypti* mosquitoes are altered by *Wolbachia* infection, which in turn alters the virome diversity [17]. The present study investigated whether differences between the microbiomes of *Ae. aegypti* and *Ae. albopictus* influence the compositions of their respective viromes. Samples collected between 2015 and the first half of 2017 that were negative for *Wolbachia* were compared with those that were positive for this bacterium. The results suggest that the presence of *w*Mel *Wolbachia* did not influence the diversity or richness of the *Ae. aegypti* virome. However, the number of *Wolbachia-*infected mosquitoes in each of the five pools of mosquitoes remained undetected. In the wild populations of *Drosophila melanogaster*, *Wolbachia* infection does not influence the diversity of native viruses in the insect [85]. Nevertheless, the results of the present study suggest that the presence of *Wolbachia* enhances the level of infection of mosquitoes by Aedes aegypti To virus 2. A similar result was observed in field-caught *Ae. aegypti* mosquitoes, where *Wolbachia* enhanced ISF infection rates and loads, demonstrating that *Wolbachia* does not act as an antiviral against all flaviviruses [24,86].

Although *Wolbachia* did not influence the *Ae. aegypti* virome diversity, a higher number of mitochondrial SNPs clustered apart from those of previous periods were observed in *Ae. aegypti* samples collected after the second half of 2017 (when the first *Wolbachia-*transfected mosquitoes were released; Fig 6). Therefore, the release of these *Wolbachia-*transfected mosquitoes affected the native *Ae. aegypti* population (Fig 7). The transfected mosquitoes were the product of a cross between the native population and *Wolbachia*-bearing mosquitoes from Australia [87]. Therefore, a temporal interpopulation substructure might have formed after transfected mosquitoes were released in Bello and Medellín. In particular, some genetic variants of the Australian strain may have remained after the crosses, and these were introduced into the Medellín mosquito population, increasing its diversity. Evidence for this can be obtained by analyzing biparental genetic markers and determining the levels of introgression between the native and introduced populations. Future *Wolbachia-*infected mosquitoes release can be facilitated to achieve faster rates of invasion at a lower cost by determining whether these variants become fixed in the native population and by assessing their effects on maintaining *Wolbachia* infections and its local spread capacities [88,89].

The analysis of mitochondrial diversity during the 3 years of *Ae. albopictus* sampling revealed lower numbers of mitochondrial variable sites and SNPs than those observed in *Ae. aegypti*; moreover, *Ae. albopictus* diversity was similar between the sampling periods. *Ae. albopictus* is an invasive species detected in Medellín in 2011 [90], and its relatively low diversity suggests that its population size is smaller than that of *Ae. aegypti*. To the best of our knowledge, however, no population genetic studies of this species have been conducted in either Medellín or Colombia. Therefore, it is not possible to establish how the diversity of this species has changed locally.

## Conclusions

In conclusion, the levels of abundance and diversity between the viromes of *Ae. aegypti* and *Ae. albopictus* differ strikingly, indicating that virome profiles are species-specific and can be determined based on their evolutionary history. Most of the viruses associated with both species were detected throughout different sampling periods, suggesting that the viromes are temporally stable. *Wolbachia* spp. from clades A and B were prevalent in *Ae. albopictus,* whereas one strain of *Wolbachia* was present in some samples of *Ae. aegypti.* The presence of *Wolbachia* in *Ae. aegypti* was not related to any changes in the composition of its virome, except for an increase in the abundance of one ISV, Aedes aegypti To virus 2. Furthermore, mitochondrial genetic diversity was lower in *Ae. albopictus* than in *Ae. aegypti*, which suggests that the population size of the former is smaller. The genetic diversity of *Ae. aegypti* increased after the second half of 2017, and this is probably related to the release of mosquitoes transfected with *Wolbachia*. This study thus provides a baseline for future virome studies on *Ae. aegypti* and *Ae. albopictus* in Colombia and other countries in the Americas.

## Acknowledgments

The authors express their sincere gratitude toward the people who allowed entry to their homes for entomological sampling as well as the Secretariat of Health of Medellín and the Medical Entomology Group of the University of Antioquia for their support during sampling.

## Author Contributions

Conceptualization, A.C.T. and A.G.P.; methodology, A.C.T., J.P.P., and A.G.P.; formal analysis A.C.T., N.F.P., and A.G.P.; investigation, A.C.T., J.P.P., O.T.C., G.R.U., and A.G.P.; writing—original draft preparation, A.C.T., and A.G.P; writing—review and editing, A.C.T., J.P.P., N.F.P., W.R.M., O.T.C., G.R.U., and A.G.P; and funding acquisition, J.P.P., G.R.U., and A.G.P.

## Data Availability Statement

The raw sequencing datasets used in the present study are available in the NCBI Sequence Read Archive repository (the Bioproject; accession code is pending). Dengue virus 3 sequence generated in this study has been deposited in GenBank (accession number is pending).

## Supplementary Material

**Figure S1**. Phylogenetic reconstruction of dengue virus sequences from *Aedes aegypti* mosquitoes in Medellín (Colombia) using a 1,057-bp region of the envelope gene. The contig was assembled from RNA-seq data, and the fragment was amplified and sequenced using the Sanger method. The sequences generated in this study are indicated in red. The scale bar indicates the number of substitutions per site.

## References

1. World Health Organization A global brief on vector-borne diseases; WHO, 2014.

2. Paupy, C.; Delatte, H.; Bagny, L.; Corbel, V.; Fontenille, D. Aedes albopictus, an arbovirus vector: From the darkness to the light. Microbes Infect. 2009, 11, 1177–1185, doi:10.1016/j.micinf.2009.05.005.

3. Ding, F.; Fu, J.; Jiang, D.; Hao, M.; Lin, G. Mapping the spatial distribution of Aedes aegypti and Aedes albopictus. Acta Trop. 2018, 178, 155–162, doi:10.1016/J.ACTATROPICA.2017.11.020.

4. Villar, L.A.; Rojas, D.P.; Besada-Lombana, S.; Sarti, E. Epidemiological Trends of Dengue Disease in Colombia (2000-2011): A Systematic Review. PLoS Negl. Trop. Dis. 2015, 9, e0003499, doi:10.1371/JOURNAL.PNTD.0003499.

5. Rico-Mendoza, A.; Alexandra, P.-R.; Chang, A.; Encinales, L.; Lynch, R. Co-circulation of dengue, chikungunya, and Zika viruses in Colombia from 2008 to 2018. Rev. Panam. Salud Pública 2019, 43, 1, doi:10.26633/RPSP.2019.49.

6. Rodriguez-Morales, A.J.; Galindo-Marquez, M.L.; Carlos, ·; García-Loaiza, J.; Juan, ·; Sabogal-Roman, A.; Marin-Loaiza, S.; Andrés, ·; Ayala, F.; Lagos-Grisales, G.J.; et al. Mapping Zika virus disease incidence in Valle del Cauca. Infection 45, doi:10.1007/s15010-016-0948-1.

7. Portilla Cabrera, C.V.; Selvaraj, J.J. Geographic shifts in the bioclimatic suitability for Aedes aegypti under climate change scenarios in Colombia. Heliyon 2020, 6, e03101, doi:10.1016/J.HELIYON.2019.E03101.

8. Singer, M. The spread of Zika and the potential for global arbovirus syndemics. *http://dx.doi.org/10.1080/17441692.2016.1225112* 2016, 12, 1–18, doi:10.1080/17441692.2016.1225112.

9. Echeverry-Cárdenas, E.; López-Castañeda, C.; Carvajal-Castro, J.D.; Aguirre-Obando, O.A. Potential geographic distribution of the tiger mosquito Aedes albopictus (Skuse, 1894) (Diptera: Culicidae) in current and future conditions for Colombia. PLoS Negl. Trop. Dis. 2021, 15, e0008212, doi:10.1371/JOURNAL.PNTD.0008212.

10. Gómez-Palacio, A.; Suaza-Vasco, J.; Castaño, S.; Triana, O.; Uribe, S. Aedes albopictus (Skuse, 1894) infected with the American-Asian genotype of dengue type 2 virus in Medellín suggests its possible role as vector of dengue fever in Colombia. Biomédica 2017, 37, 135, doi:10.7705/biomedica.v37i0.3474.

11. Calle-Tobón, A.; Pérez-Pérez, J.; Rojo, R.; Rojas-Montoya, W.; Triana-Chavez, O.; Rúa-Uribe, G.; Gómez-Palacio, A. Surveillance of Zika virus in field-caught Aedes aegypti and Aedes albopictus suggests important role of male mosquitoes in viral populations maintenance in Medellín, Colombia. Infect. Genet. Evol. 2020, 85, 104434, doi:10.1016/J.MEEGID.2020.104434.

12. Secretaría de Salud de Medellín 2010 Boletin epidemiológico semana 52; 2010;

13. Secretaría de salud de Medellín Informe de periodo epidemiológico Medellín 2016. 2016, 52, 91.

14. Walker, T.; Johnson, P.H.; Moreira, L.A.; Iturbe-Ormaetxe, I.; Frentiu, F.D.; McMeniman, C.J.; Leong, Y.S.; Dong, Y.; Axford, J.; Kriesner, P.; et al. The wMel Wolbachia strain blocks dengue and invades caged Aedes aegypti populations. Nat. 2011 4767361 2011, 476, 450–453, doi:10.1038/nature10355.

15. Caragata, E.P.; Dutra, H.L.C.; Moreira, L.A. Inhibition of Zika virus by Wolbachia inAedes aegypti. Microb. Cell 2016, 3, 293, doi:10.15698/MIC2016.07.513.

16. Hurk, A.F. van den; Hall-Mendelin, S.; Pyke, A.T.; Frentiu, F.D.; McElroy, K.; Day, A.; Higgs, S.; O’Neill, S.L. Impact of Wolbachia on Infection with Chikungunya and Yellow Fever Viruses in the Mosquito Vector Aedes aegypti. PLoS Negl. Trop. Dis. 2012, 6, e1892, doi:10.1371/JOURNAL.PNTD.0001892.

17. Audsley, M.D.; Seleznev, A.; Joubert, D.A.; Woolfit, M.; O’Neill, S.L.; McGraw, E.A. Wolbachia infection alters the relative abundance of resident bacteria in adult Aedes aegypti mosquitoes, but not larvae. Mol. Ecol. 2018, 27, 297–309, doi:10.1111/MEC.14436.

18. Feschotte, C.; Gilbert, C. Endogenous viruses: insights into viral evolution and impact on host biology. Nat. Rev. Genet. 2012 134 2012, 13, 283–296, doi:10.1038/nrg3199.

19. Guégan, M.; Zouache, K.; Démichel, C.; Minard, G.; Tran Van, V.; Potier, P.; Mavingui, P.; Valiente Moro, C. The mosquito holobiont: fresh insight into mosquito-microbiota interactions. Microbiome 2018 61 2018, 6, 1–17, doi:10.1186/S40168-018-0435-2.

20. Du, J.; Li, F.; Han, Y.; Fu, S.; Liu, B.; Shao, N.; Su, H.; Zhang, W.; Zheng, D.; Lei, W.; et al. Characterization of viromes within mosquito species in China. Sci. China Life Sci. 2019, 63, 1089–1092, doi:10.1007/s11427-019-1583-9.

21. Nanfack-Minkeu, F.; Mitri, C.; Bischoff, E.; Belda, E.; Casademont, I.; Vernick, K.D. Interaction of RNA viruses of the natural virome with the African malaria vector, Anopheles coluzzii. Sci. Rep. 2019, 9, 1–10, doi:10.1038/s41598-019-42825-3.

22. Shi, C.; Zhao, L.; Atoni, E.; Zeng, W.; Hu, X.; Matthijnssens, J.; Yuan, Z.; Xia, H. Stability of the Virome in Lab- and Field-Collected Aedes albopictus Mosquitoes across Different Developmental Stages and Possible Core Viruses in the Publicly Available Virome Data of Aedes Mosquitoes. mSystems 2020, 5, 1–16, doi:10.1128/msystems.00640-20.

23. Xiao, P.; Han, J.; Zhang, Y.; Li, C.; Guo, X.; Wen, S.; Tian, M.; Li, Y.; Wang, M.; Liu, H.; et al. Metagenomic Analysis of Flaviviridae in Mosquito Viromes Isolated From Yunnan Province in China Reveals Genes From Dengue and Zika Viruses. Front. Cell. Infect. Microbiol. 2018, 8, 359, doi:10.3389/fcimb.2018.00359.

24. Zakrzewski, M.; Rašić, G.; Darbro, J.; Krause, L.; Poo, Y.S.; Filipović, I.; Parry, R.; Asgari, S.; Devine, G.; Suhrbier, A. Mapping the virome in wild-caught Aedes aegypti from Cairns and Bangkok. Sci. Rep. 2018, 8, 1–12, doi:10.1038/s41598-018-22945-y.

25. Sadeghi, M.; Altan, E.; Deng, X.; Barker, C.M.; Fang, Y.; Coffey, L.L.; Delwart, E. Virome of > 12 thousand Culex mosquitoes from throughout California. Virology 2018, 523, 74–88, doi:10.1016/j.virol.2018.07.029.

26. Shi, C.; Zhao, L.; Atoni, E.; Zeng, W.; Hu, X.; Matthijnssens, J.; Yuan, Z.; Xia, H. The conservation of a core virome in Aedes mosquitoes across different developmental stages and continents. bioRxiv 2020, 2020.04.23.058701, doi:10.1101/2020.04.23.058701.

27. Nouri, S.; Matsumura, E.E.; Kuo, Y.W.; Falk, B.W. Insect-specific viruses: from discovery to potential translational applications. Curr. Opin. Virol. 2018, 33, 33–41, doi:10.1016/J.COVIRO.2018.07.006.

28. Patterson, E.I.; Villinger, J.; Muthoni, J.N.; Dobel-Ober, L.; Hughes, G.L. Exploiting insect-specific viruses as a novel strategy to control vector-borne disease. Curr. Opin. Insect Sci. 2020, 39, 50–56, doi:10.1016/J.COIS.2020.02.005.

29. Öhlund, P.; Lundén, H.; Blomström, A.-L. Insect-specific virus evolution and potential effects on vector competence. Virus Genes 2019 552 2019, 55, 127–137, doi:10.1007/S11262-018-01629-9.

30. Romo, H.; Kenney, J.L.; Blitvich, B.J.; Brault, A.C. Restriction of Zika virus infection and transmission in Aedes aegypti mediated by an insect-specific flavivirus. Emerg. Microbes Infect. 2018, 7, doi:10.1038/s41426-018-0180-4.

31. Olmo, R.P.; Todjro, Y.M.H.; Aguiar, E.R.G.R.; Almeida, J.P.P. de; Armache, J.N.; Faria, I.J.S. de; Ferreira, F. V.; Silva, A.T.S.; Souza, K.P.R. de; Vilela, A.P.P.; et al. Insect-specific viruses regulate vector competence in Aedes aegypti mosquitoes via expression of histone H4. bioRxiv 2021, 2021.06.05.447047, doi:10.1101/2021.06.05.447047.

32. Pérez-Pérez, J.; Sanabria, W.H.; Restrepo, C.; Rojo, R.; Henao, E.; Triana, O.; Mejía, A.M.; Castaño, S.M.; Rúa-Uribe, G.L. Virological surveillance of Aedes (Stegomyia) aegypti and Aedes (Stegomyia) albopictus as support for decision making for dengue control in Medellín. Biomédica 2017, 37, 155, doi:10.7705/biomedica.v37i0.3467.

33. Pérez-Pérez, J.; Rojo, R.; Henao, E.; García-Huerta, P.; Triana-Chavez, O.; Rúa-Uribe, G. Natural infection of Aedes aegypti, Ae. albopictus and Culex spp. with Zika virus in Medellin, Colombia. CES Med. 2019, 33, 175–181, doi:10.21615/cesmedicina.33.3.2.

34. Velez, I.D.; Santacruz, E.; Kutcher, S.C.; Duque, S.L.; Uribe, A.; Barajas, J.; Gonzalez, S.; Patino, A.C.; Zuluaga, L.; Martínez, L.; et al. The impact of city-wide deployment of Wolbachia-carrying mosquitoes on arboviral disease incidence in Medellín and Bello, Colombia: study protocol for an interrupted time-series analysis and a test-negative design study. F1000Research 2019, 8, 1327, doi:10.12688/F1000RESEARCH.19858.1.

35. Rueda, L.M. Pictorial keys for the identification of mosquitoes (Diptera: Culicidae) associated with Dengue Virus Transmission. Zootaxa 2004, 589, 1, doi:10.11646/zootaxa.589.1.1.

36. Chen, S.; Zhou, Y.; Chen, Y.; Gu, J. fastp: an ultra-fast all-in-one FASTQ preprocessor. Bioinformatics 2018, 34, i884–i890, doi:10.1093/bioinformatics/bty560.

37. Matthews, B.J.; Dudchenko, O.; Kingan, S.B.; Koren, S.; Antoshechkin, I.; Crawford, J.E.; Glassford, W.J.; Herre, M.; Redmond, S.N.; Rose, N.H.; et al. Improved reference genome of Aedes aegypti informs arbovirus vector control. Nat. 2018 5637732 2018, 563, 501–507, doi:10.1038/s41586-018-0692-z.

38. Chen, X.-G.; Jiang, X.; Gu, J.; Xu, M.; Wu, Y.; Deng, Y.; Zhang, C.; Bonizzoni, M.; Dermauw, W.; Vontas, J.; et al. Genome sequence of the Asian Tiger mosquito, Aedes albopictus, reveals insights into its biology, genetics, and evolution. Proc. Natl. Acad. Sci. 2015, 112, E5907–E5915, doi:10.1073/PNAS.1516410112.

39. Li, H.; Durbin, R. Fast and accurate short read alignment with Burrows-Wheeler transform. Bioinformatics 2009, 25, 1754–1760, doi:10.1093/bioinformatics/btp324.

40. Kopylova, E.; Noé, L.; Touzet, H. SortMeRNA: fast and accurate filtering of ribosomal RNAs in metatranscriptomic data. Bioinformatics 2012, 28, 3211–3217, doi:10.1093/bioinformatics/bts611.

41. Zhao, L.; Atoni, E.; Shi, C.; Yuan, Z.; Xia, H. Mapping the virome in lab-reared and wild-caught Aedes albopictus mosquitoes. Access Microbiol. 2019, 1, 4, doi:10.1099/acmi.imav2019.po0009.

42. Bankevich, A.; Nurk, S.; Antipov, D.; Gurevich, A.A.; Dvorkin, M.; Kulikov, A.S.; Lesin, V.M.; Nikolenko, S.I.; Pham, S.; Prjibelski, A.D.; et al. SPAdes: A new genome assembly algorithm and its applications to single-cell sequencing. J. Comput. Biol. 2012, 19, 455–477, doi:10.1089/cmb.2012.0021.

43. Buchfink, B.; Xie, C.; Huson, D.H. Fast and sensitive protein alignment using DIAMOND. Nat. Methods 2014, 12, 59–60.

44. Kubacki, J.; Flacio, E.; Qi, W.; Guidi, V.; Tonolla, M.; Fraefel, C. Viral Metagenomic Analysis of Aedes albopictus Mosquitos from Southern Switzerland. Viruses 2020, 12, 929, doi:10.3390/v12090929.

45. Ondov, B.D.; Bergman, N.H.; Phillippy, A.M. Interactive metagenomic visualization in a Web browser. BMC Bioinformatics 2011, 12, doi:10.1186/1471-2105-12-385.

46. Lagkouvardos, I.; Fischer, S.; Kumar, N.; Clavel, T. Rhea: a transparent and modular R pipeline for microbial profiling based on 16S rRNA gene amplicons. PeerJ 2017, 5, e2836, doi:10.7717/PEERJ.2836.

47. R Core Team R: A language and environment for statistical computing 2020.

48. Anderson, M.J.; Crist, T.O.; Chase, J.M.; Vellend, M.; Inouye, B.D.; Freestone, A.L.; Sanders, N.J.; Cornell, H. V.; Comita, L.S.; Davies, K.F.; et al. Navigating the multiple meanings of β diversity: a roadmap for the practicing ecologist. Ecol. Lett. 2011, 14, 19–28, doi:10.1111/J.1461-0248.2010.01552.X.

49. Li, H. Aligning sequence reads, clone sequences and assembly contigs with BWA-MEM. 2013.

50. h, L.; b, H.; a, W.; t, F.; j, R.; n, H.; g, M.; g, A.; r, D. The Sequence Alignment/Map format and SAMtools. Bioinformatics 2009, 25, 2078–2079, doi:10.1093/BIOINFORMATICS/BTP352.

51. h, L. A statistical framework for SNP calling, mutation discovery, association mapping and population genetical parameter estimation from sequencing data. Bioinformatics 2011, 27, 2987–2993, doi:10.1093/BIOINFORMATICS/BTR509.

52. Garrison, E.; Marth, G. Haplotype-based variant detection from short-read sequencing. 2012.

53. DePristo, M.A.; Banks, E.; Poplin, R.; Garimella, K. V; Maguire, J.R.; Hartl, C.; Philippakis, A.A.; del Angel, G.; Rivas, M.A.; Hanna, M.; et al. A framework for variation discovery and genotyping using next-generation DNA sequencing data. Nat. Genet. 2011 435 2011, 43, 491–498, doi:10.1038/ng.806.

54. Rozas, J.; Ferrer-Mata, A.; Sánchez-DelBarrio, J.C.; Guirao-Rico, S.; Librado, P.; Ramos-Onsins, S.E.; Sánchez-Gracia, A. DnaSP 6: DNA Sequence Polymorphism Analysis of Large Data Sets. Mol. Biol. Evol. 2017, 34, 3299–3302, doi:10.1093/molbev/msx248.

55. Kumar, S.; Stecher, G.; Li, M.; Knyaz, C.; Tamura, K. MEGA X: Molecular Evolutionary Genetics Analysis across Computing Platforms. Mol. Biol. Evol. 2018, 35, 1547, doi:10.1093/MOLBEV/MSY096.

56. Jiménez-Silva, C.L.; Carreño, M.F.; Ortiz-Baez, A.S.; Rey, L.A.; Villabona-Arenas, C.J.; Ocazionez, R.E. Evolutionary history and spatio-temporal dynamics of dengue virus serotypes in an endemic region of Colombia. PLoS One 2018, 13, e0203090, doi:10.1371/journal.pone.0203090.

57. Hall, T. BioEdit v.7.0.5. Biological Sequences Alignment Editor for Windows. Ibis Therapeutics, a Division of Isis Pharmaceuticals. 2005.

58. Katoh, K.; Standley, D.M. MAFFT Multiple Sequence Alignment Software Version 7: Improvements in Performance and Usability. Mol. Biol. Evol. 2013, 30, 772–780, doi:10.1093/molbev/mst010.

59. Nguyen, L.-T.; Schmidt, H.A.; von Haeseler, A.; Minh, B.Q. IQ-TREE: A Fast and Effective Stochastic Algorithm for Estimating Maximum-Likelihood Phylogenies. Mol. Biol. Evol. 2015, 32, 268–274, doi:10.1093/molbev/msu300.

60. Kalyaanamoorthy, S.; Minh, B.Q.; Wong, T.K.F.; Von Haeseler, A.; Jermiin, L.S. ModelFinder: Fast model selection for accurate phylogenetic estimates. Nat. Methods 2017, 14, 587–589, doi:10.1038/nmeth.4285.

61. Shi, C.; Beller, L.; Deboutte, W.; Yinda, K.C.; Delang, L.; Vega-Rúa, A.; Failloux, A.B.; Matthijnssens, J. Stable distinct core eukaryotic viromes in different mosquito species from Guadeloupe, using single mosquito viral metagenomics. Microbiome 2019, 7, 1–20, doi:10.1186/s40168-019-0734-2.

62. x, Z.; s, H.; t, J.; p, L.; y, H.; c, W.; b, P.; l, W.; h, C.; m, W.; et al. Discovery and high prevalence of Phasi Charoen-like virus in field-captured Aedes aegypti in South China. Virology 2018, 523, 35–40, doi:10.1016/J.VIROL.2018.07.021.

63. Chandler, J.A.; Thongsripong, P.; Green, A.; Kittayapong, P.; Wilcox, B.A.; Schroth, G.P.; Kapan, D.D.; Bennett, S.N. Metagenomic shotgun sequencing of a Bunyavirus in wild-caught Aedes aegypti from Thailand informs the evolutionary and genomic history of the Phleboviruses. Virology 2014, *464*–*465*, 312–319, doi:10.1016/J.VIROL.2014.06.036.

64. Ramos-Nino, M.E.; Fitzpatrick, D.M.; Tighe, S.; Eckstrom, K.M.; Hattaway, L.M.; Hsueh, A.N.; Stone, D.M.; Dragon, J.; Cheetham, S. High prevalence of Phasi Charoen-like virus from wild-caught Aedes aegypti in Grenada, W.I. as revealed by metagenomic analysis. PLoS One 2020, 15, e0227998, doi:10.1371/journal.pone.0227998.

65. Aubry, F.; Dabo, S.; Manet, C.; Filipović, I.; Rose, N.H.; Miot, E.F.; Martynow, D.; Baidaliuk, A.; Merkling, S.H.; Dickson, L.B.; et al. Enhanced Zika virus susceptibility of globally invasive Aedes aegypti populations. Science (80-.). 2020, 370, 991–996, doi:10.1126/SCIENCE.ABD3663.

66. de Oliveira Ribeiro, G.; Morais, V.S.; Monteiro, F.J.C.; Ribeiro, E.S.D.A.; da Rego, M.O.S.; Souto, R.N.P.; Villanova, F.; Tahmasebi, R.; Hefford, P.M.; Deng, X.; et al. Aedes aegypti from amazon basin harbor high diversity of novel viral species. Viruses 2020, 12, doi:10.3390/v12080866.

67. Stollar, V.; Thomas, V.L. An agent in the Aedes aegypti cell line (Peleg) which causes fusion of Aedes albopictus cells. Virology 1975, 64, 367–377, doi:10.1016/0042-6822(75)90113-0.

68. Espinoza-Gómez, F.; López-Lemus, A.U.; Rodriguez-Sanchez, I.P.; Martinez-Fierro, M.L.; Newton-Sánchez, O.A.; Chávez-Flores, E.; Delgado-Enciso, I. Detection of sequences from a potentially novel strain of cell fusing agent virus in Mexican Stegomyia (Aedes) aegypti mosquitoes., doi:10.1007/s00705-011-0967-2.

69. Cook, S.; Bennett, S.N.; Holmes, E.C.; Chesse, R. De; Moureau, G.; Lamballerie, X. de Isolation of a new strain of the flavivirus cell fusing agent virus in a natural mosquito population from Puerto Rico. J. Gen. Virol. 2006, 87, 735–748, doi:10.1099/VIR.0.81475-0.

70. Yamanaka, A.; Thongrungkiat, S.; Ramasoota, P.; Konishi, E. Genetic and evolutionary analysis of cell-fusing agent virus based on Thai strains isolated in 2008 and 2012. Infect. Genet. Evol. 2013, 19, 188–194, doi:10.1016/J.MEEGID.2013.07.012.

71. Zhang, G.; Asad, S.; Khromykh, A.A.; Asgari, S. Cell fusing agent virus and dengue virus mutually interact in Aedes aegypti cell lines. Sci. Reports 2017 71 2017, 7, 1–8, doi:10.1038/s41598-017-07279-5.

72. Zhang, X.; Guo, • Xiaofang; Fan, • Hang; Zhao, Q.; Zuo, S.; Qiang Sun, •; Pei, G.; Cheng, • Shi; An, X.; Wang, Y.; et al. Complete genome sequence of Menghai flavivirus, a novel insect-specific flavivirus from China. Arch. Virol. 162, doi:10.1007/s00705-017-3232-5.

73. Parry, R.; Naccache, F.; Ndiaye, E.H.; Fall, G.; Castelli, I.; Lühken, R.; Medlock, J.; Cull, B.; Hesson, J.C.; Montarsi, F.; et al. Identification and RNAi Profile of a Novel Iflavirus Infecting Senegalese Aedes vexans arabiensis Mosquitoes. Viruses 2020, Vol. 12, Page 440 2020, 12, 440, doi:10.3390/V12040440.

74. Kobayashi, D.; Isawa, H.; Fujita, R.; Murota, K.; Itokawa, K.; Higa, Y.; Katayama, Y.; Sasaki, T.; Mizutani, T.; Iwanaga, S.; et al. Isolation and characterization of a new iflavirus from Armigeres spp. mosquitoes in the Philippines. J. Gen. Virol. 2017, 98, 2876–2881, doi:10.1099/JGV.0.000929.

75. Parry, R.; Asgari, S. Aedes Anphevirus: an Insect-Specific Virus Distributed Worldwide in Aedes aegypti Mosquitoes That Has Complex Interplays with Wolbachia and Dengue Virus Infection in Cells. J. Virol. 2018, 92, 1–19, doi:10.1128/jvi.00224-18.

76. Hoshino, K.; Isawa, H.; Tsuda, Y.; Sawabe, K.; Kobayashi, M. Isolation and characterization of a new insect flavivirus from Aedes albopictus and Aedes flavopictus mosquitoes in Japan. Virology 2009, 391, 119–129, doi:10.1016/J.VIROL.2009.06.025.

77. Fang, Y.; Zhang, Y.; Zhou, Z.-B.; Shi, W.-Q.; Xia, S.; Li, Y.-Y.; Wu, J.-T.; Liu, Q.; Lin, G.-Y. Co-circulation of Aedes flavivirus, Culex flavivirus, and Quang Binh virus in Shanghai, China. Infect. Dis. Poverty 2018 71 2018, 7, 1–10, doi:10.1186/S40249-018-0457-9.

78. Schultz, M.J.; Frydman, H.M.; Connor, J.H. Dual Insect specific virus infection limits Arbovirus replication in Aedes mosquito cells. Virology 2018, 518, 406–413, doi:10.1016/J.VIROL.2018.03.022.

79. Gratz, N.G. Critical review of the vector status of Aedes albopictus. Med. Vet. Entomol. 2004, 18, 215–227, doi:10.1111/j.0269-283X.2004.00513.x.

80. Richards, S.L.; Ponnusamy, L.; Unnasch, T.R.; Hassan, H.K.; Apperson, C.S. Host-Feeding Patterns of *Aedes albopictus* (Diptera: Culicidae) in Relation to Availability of Human and Domestic Animals in Suburban Landscapes of Central North Carolina. J. Med. Entomol. 2006, 43, 543–551, doi:10.1093/jmedent/43.3.543.

81. Wang, L.; Lv, X.; Zhai, Y.; Fu, S.; Wang, D.; Rayner, S.; Tang, Q.; Liang, G. Genomic Characterization of a Novel Virus of the Family Tymoviridae Isolated from Mosquitoes. PLoS One 2012, 7, e39845, doi:10.1371/JOURNAL.PONE.0039845.

82. Ahmad, N.A.; Vythilingam, I.; Lim, Y.A.L.; Zabari, N.Z.A.M.; Lee, H.L. Detection of Wolbachia in aedes albopictus and their effects on chikungunya virus. Am. J. Trop. Med. Hyg. 2017, 96, 148–156, doi:10.4269/ajtmh.16-0516.

83. Seabournid, P.; Spafford, H.; Yoneishi, N.; Medeirosid, M. The aedes albopictus (Diptera: Culicidae) microbiome varies spatially and with ascogregarine infection. PLoS Negl. Trop. Dis. 2020, 14, 1–21, doi:10.1371/journal.pntd.0008615.

84. Ross, P.A.; Callahan, A.G.; Yang, Q.; Jasper, M.; Arif, M.A.K.; Afizah, A.N.; Nazni, W.A.; Hoffmann, A.A. An elusive endosymbiont: Does Wolbachia occur naturally in Aedes aegypti? Ecol. Evol. 2020, 10, 1581–1591, doi:10.1002/ECE3.6012.

85. Shi, M.; White, V.L.; Schlub, T.; Eden, J.S.; Hoffmann, A.A.; Holmes, E.C. No detectable effect of Wolbachia wMel on the prevalence and abundance of the RNA virome of Drosophila melanogaster. Proc. R. Soc. B Biol. Sci. 2018, 285, doi:10.1098/rspb.2018.1165.

86. Amuzu, H.E.; Tsyganov, K.; Koh, C.; Herbert, R.I.; Powell, D.R.; McGraw, E.A. Wolbachia enhances insect-specific flavivirus infection in Aedes aegypti mosquitoes. Ecol. Evol. 2018, 8, 5441–5454, doi:10.1002/ece3.4066.

87. Aliota, M.T.; Peinado, S.A.; Velez, I.D.; Osorio, J.E. The wMel strain of Wolbachia Reduces Transmission of Zika virus by Aedes aegypti. Sci. Reports 2016 61 2016, 6, 1–7, doi:10.1038/srep28792.

88. Schmidt, T.L.; Filipović, I.; Hoffmann, A.A.; Rašić, G. Fine-scale landscape genomics helps explain the slow spatial spread of Wolbachia through the Aedes aegypti population in Cairns, Australia. Hered. 2018 1205 2018, 120, 386–395, doi:10.1038/s41437-017-0039-9.

89. Jasper, M.; Schmidt, T.L.; Ahmad, N.W.; Sinkins, S.P.; Hoffmann, A.A. A genomic approach to inferring kinship reveals limited intergenerational dispersal in the yellow fever mosquito. Mol. Ecol. Resour. 2019, 19, 1254–1264, doi:10.1111/1755-0998.13043.

90. Rúa-Uribe, G.; Suárez Acosta, C.; Londoño, V.; Sanchez, J.; Rojo, R.; Novoa, B. Primera evidencia de Aedes albopictus (Skuse) (Diptera: Culicidae) en la ciudad de Medellín, Antioquia - Colombia. Rev. Salud Pública Medellín 2011, 5, 89–98.

